# Identification of distinct subtypes of post-stroke and neurotypical gait behaviors using neural network analysis of kinematic time series data

**DOI:** 10.1101/2024.10.28.620665

**Authors:** Andrian Kuch, Nicolas Schweighofer, James M. Finley, Alison McKenzie, Yuxin Wen, Natalia Sánchez

**Affiliations:** Department of Physical Therapy, Chapman University, Irvine, CA; Division of Biokinesiology and Physical Therapy; Department of Biomedical Engineering; Neuroscience Graduate Program, University of Southern California, Los Angeles, CA; Fowler School of Engineering, Chapman University, Orange, CA

**Keywords:** Stroke, gait, rehabilitation, machine learning, neural networks, clustering, kinematics

## Abstract

**Background:** Heterogeneous types of gait impairment are common post-stroke. Studies used supervised and unsupervised machine learning on discrete biomechanical features to summarize the gait cycle and identify common patterns of gait behaviors. However, discrete features cannot account for temporal variations that occur during gait. Here, we propose a novel machine-learning pipeline to identify subgroups of gait behaviors post-stroke using kinematic time series data.

**Methods:** We analyzed ankle and knee kinematic data during treadmill walking data in 39 individuals post-stroke and 28 neurotypical controls. The data were first input into a supervised dual-stage Convolutional Neural Network-Temporal Convolutional Network, trained to extract temporal and spatial gait features. Then, we used these features to find clusters of different gait behaviors using unsupervised time series k-means. We repeated the clustering process using 10,000 bootstrap training data samples and a Gaussian Mixture Model to identify stable clusters representative of our dataset. Finally, we assessed the kinematic differences between the identified clusters using 1D statistical parametric mapping ANOVA. We then compared gait spatiotemporal and clinical characteristics between clusters using one-way ANOVA.

**Results:** We obtained five clusters: two clusters of neurotypical individuals (C1 and C2) and three clusters of individuals post-stroke (S1, S2, S3). C1 had kinematics that resembled the normative gait pattern. Individuals in C2 had a shorter stride time than C1. Individuals in S1 had mild impairment and walked with increased bilateral knee flexion during the loading response. Individuals in S2 had moderate impairment, were the slowest among the clusters, took shorter steps, had increased knee flexion during stance bilaterally and reduced paretic knee flexion during swing. Individuals in S3 had mild impairment, asymmetric swing time, had increased ankle abduction during the gait cycle and reduced dorsiflexion bilaterally during loading response and stance. Every individual was assigned to a cluster with a cluster membership likelihood above 93%.

**Conclusions:** Our results indicate that joint kinematics in individuals post-stroke are distinct from controls, even in those individuals with mild impairment. The three subgroups post-stroke showed distinct kinematic impairments during specific phases in the gait cycle, providing additional information to clinicians for gait retraining interventions.

## Background

Gait patterns differ between stroke survivors due to heterogeneity in stroke lesion type, size, location, and differences in recovery ^1–7^. These differences make intervention prescription difficult in both research and rehabilitation interventions. Developing approaches to identify intervention targets systematically can enhance the efficacy of physical therapy interventions to improve walking function in stroke survivors.

The different types of gait patterns post-stroke have been identified qualitatively and quantitatively in prior research studies ^1,2,5,6,8^. Using visual assessment of paretic electromyography (EMG), a seminal study ^1^ identified three subgroups of abnormal muscle activation during gait post-stroke based on activation onset and levels – early triceps surae activation, decreased activation of paretic musculature, and paretic muscle coactivation ^1^. Similarly, Olney and Richards qualitatively identified different subgroups of gait impairments using peak spatiotemporal, peak kinematic, and peak kinetic characteristics ^6^. A more systematic quantitative approach used paretic EMGs (onset and percentages of maximum voluntary contraction) and paretic peak kinematics input into a hierarchical clustering algorithm to identify four clusters of gait behaviors post-stroke ^2^: a fast walking group with slight knee flexion in mid-stance, an intermediate velocity group with increased knee flexion in mid-stance, a slow group with excessive knee flexion in midstance and a slow group with knee hyperextension in mid-stance ^2^. Our recent work used spatiotemporal variables and peak ground reaction forces from both the paretic and non-paretic extremity input into a k-means clustering algorithm to identify four types of gait behaviors in individuals post-stroke ^5^: fast and asymmetric walkers, moderate speed, and asymmetric walkers, slow walkers with frontal plane impairment, and slow and symmetric walkers. While all these previous studies have provided valuable information to identify different types of gait behaviors post-stroke, a caveat is that they have used discrete metrics over the gait cycle ^2,5,6,8^. These discrete gait metrics summarize changes in magnitude but not in the timing of gait kinematics, kinetics, and EMGs that result in gait impairment post-stroke. Thus, these previous studies cannot identify specific phases of the gait cycle that could be targeted in rehabilitation interventions.

The use of machine learning methods to study gait has become common practice in research ^9^. Previous studies using supervised methods aimed to accurately classify neurotypical and pathological gait or detect types of activities ^9^. Most of these studies use handpicked discrete summary features ^9^. However, machine learning algorithms can be leveraged to select more complex and objective features using multivariate time series. Cui et al. ^10^ proposed a framework using neural networks to handle multidimensional time series to classify neurotypical and post-stroke gait and assess walking quality. While providing a walking quality score is promising, it does not inform clinicians what to target systematically during rehabilitation interventions. Unsupervised methods, on the other hand can be employed to identify clusters in a dataset ^11^. For example, Pulido-Valdeolivas et al ^12^ used time series and combined dynamic time-warping algorithms with unsupervised clustering methods to identify clusters of gait behaviors in individuals with hereditary spastic paraplegia. The dynamic time warping approach of the previous study can handle multivariate time series and uses a distance metric to compare the similarity between signals that may differ in duration ^13^. However, a completely unsupervised approach can be sensitive to outliers dependent on the choice of distance measure and leading to results that are not always generalizable, thus difficult to interpret ^11^. Therefore, using a first supervised layer to accurately extract features distinct between individuals post-stroke and neurotypical controls before a second unsupervised machine learning methods to identify the clusters might be a more suitable approach to identify clusters of gait behaviors post-stroke.

Here, we designed a pipeline to analyze time series data that combines supervised and unsupervised analyses. First, we use supervised analyses to extract features from time series data that best distinguish between individuals post-stroke and neurotypical controls. Then, we use unsupervised clustering using features weights of the supervised stage to identify clusters of gait behaviors. We compared the performance of our proposed pipeline with an unsupervised time series clustering algorithm available in R ^13^. We developed and validated our approach using kinematic data, as these data can be assessed in the clinic via visual gait analysis or simple video gait analysis ^14–16^. We implemented our pipeline in a sample of participants post-stroke and age-matched controls, to determine whether less impaired individuals post-stroke could be comparable to neurotypical controls. We hypothesized that we would observe distinct clusters of gait behaviors in individuals post-stroke ^1,2,5^, and a cluster of control individuals and individuals post-stroke with gait behaviors indistinguishable from controls ^5^, indicative of full recovery of gait post-stroke. Our proposed pipeline can be applied to other time series data during different motor tasks and to other pathological populations to identify subtypes of behaviors across different variables and populations.

## Methods

Data for a total of = 67 participants, including 39 individuals post-stroke and 28 age and sex-matched controls were curated from previous studies (Table 1) ^5,17,18^. Inclusion criteria for participants post-stroke were: (1) a unilateral stroke more than six months before the study, (2) paresis confined to one side, (3) ability to provide informed consent and communicate with the investigators, and (4) ability to walk 5 minutes on a treadmill without the assistance of another individual or walking aids (e.g., a cane or walker). The use of an ankle-foot orthosis or brace was permitted during data collection. Inclusion criteria for neurotypical participants were: (1) being of the same age and sex as a participant post-stroke, (2) having no musculoskeletal or neurologic injury that hinders walking ability, (3) ability to provide informed consent and communicate with the investigators.

**Table 1:** Participant demographics. Descriptive statistics are presented as average ± standard deviation with the range in brackets. F: Female, M: Male, SS: Self-selected, FM: lower extremity Fugl-Meyer score, ABC: Activities Balance Confidence Scale, FGA: Functional Gait Assessment, BBS: Berg Balance Score, L: Left, R: Right. *Significant differences between participants post-stroke and controls (p<0.05)

|  | Stroke | Control |
| --- | --- | --- |
| **N** | 39 | 28 |
| **Sex** | 17F/22M | 16F/12M |
| **Age (years)** | 59.5 ± 10.8 [29 – 78] | 62.4 ± 14.2 [24 – 81] |
| **Mass (kg)** | 74.1 ± 16.3 [45 – 104] | 73 ± 15.7 [46 – 110] |
| **Height (m)** | 1.58 ± 0.08 [1.44 – 1.80] | 1.60 ± 0.09 [1.42 – 1.74] |
| **Treadmill Speed (m/s)** | 0.56 ± 0.20 [0.20 – 0.95] | SS: 0.84 ± 0.26 [0.48 – 1.43] |
| **ABC (100 max)** | 74.0 ± 18.0* [38 – 98] | 95.7 ± 4.7 [83.75 – 100] |
| **BBS (56 max)** | 50.8 ± 5.5* [49 – 56] | 54.1 ± 2.6 [49 – 56] |
| **FM (34 max)** | 26.6 ± 4.83 [15 – 33] |  |
| **FGA (30 max)** | 21.2 ± 0.95 [6 - 30] |  |
| **Paresis** | 22R/17L |  |
| **Time post-stroke** | 92 ± 84.5 [6 – 467] months |  |

### Experimental Protocol for Data Collection

We performed the following assessments in both participants post-stroke and neurotypical controls: Berg Balance Scale (BBS) ^19^, Activity-Specific Balance Confidence (ABC) test ^20^, and 10-meter walk test. In participants post-stroke, we performed the lower extremity motor domain of the Fugl-Meyer (FM) assessment of motor impairment ^21^ and the Functional Gait Assessment (FGA) ^22^.

After clinical assessments, we determined participants’ self-selected speed on an instrumented treadmill (Bertec, Colombus, USA) using the staircase method ^23^, by progressively increasing or decreasing the speed in increments of 0.05 m/s until the participant felt that the speed was comfortable. The treadmill speed was required to be at least 70% of their overground walking speed measured via the 10-meter walk test. Post-stroke participants walked at this speed for three minutes, instructed to walk as it felt natural. For neurotypical controls, participants walked at their self-selected speed and the speed of a participant post-stroke matched for age and sex. All clustering analyses in control participants used data collected while walking at a matched speed of a stroke participant of the same age and sex to differentiate impairments due to stroke from those due to walking at a slower speed.

Segmental kinematics were recorded using a full-body marker set, placing retroreflective markers on bony landmarks and marker clusters over the upper arms, lower arms, thighs, shanks, and heels ^24,25^. Marker data were recorded using a 10-camera Qualisys Oqus motion capture system (Qualisys AB, Göteborg, Sweden) at 100 Hz. Forces were measured from force plates embedded in the treadmill at a sample 1000 Hz.

### Data Processing

Markers were labeled using Qualisys Track Manager and then exported to Visual 3D (C-Motion, Kingston, Canada) to construct a full-body model. Three-dimensional marker positions were filtered using a Butterworth lowpass filter with a 6 Hz cutoff frequency. We then created a body model in Visual 3D with the following segments: trunk, thighs, shanks, and feet. Participants post-stroke wore a safety harness over their pelvis to prevent falls ^18^. Thus, we removed the reflective markers from the pelvis, which prevented us from calculating hip and pelvis kinematics. All data were exported from Visual 3D to MATLAB for further processing.

Data processing and analysis were done in MATLAB (2023b, The MathWorks Inc, Natick, USA) using custom-written code. Kinematic data for the ankle in the plantar/dorsiflexion degree of freedom, ankle abduction/adduction degrees of freedom, and the knee in the flexion/extension degrees of freedom were extracted for the middle 50 seconds of the walking trial for both extremities in all participants (Fig. 1 A). Data were segmented into strides using ground reaction forces, with a threshold of 32 N ^26,27^ to detect initial contact. All strides were interpolated in time to 101 samples, with the first sample corresponding to 0% of the gait cycle or initial contact and 100% corresponding to the end of the gait cycle. We obtained the median over the gait cycle across all strides to reduce the influence of outlier stepping patterns, thus obtaining a representative stride for each degree of freedom in both extremities for each participant (Fig. 1 A). In participants post-stroke, we identified the paretic and non-paretic extremities. For data collected in neurotypical adults, leg dominance was defined as the leg they would use to kick a ball, which was the right leg for all participants. Control data are labeled as dominant and non-dominant, and comparisons are made for the non-dominant leg vs. the paretic leg, and the dominant leg vs. the non-paretic leg. 0% of the gait cycle is expressed relative to the non-dominant or paretic heel strike for both limbs.

**Figure 1:**
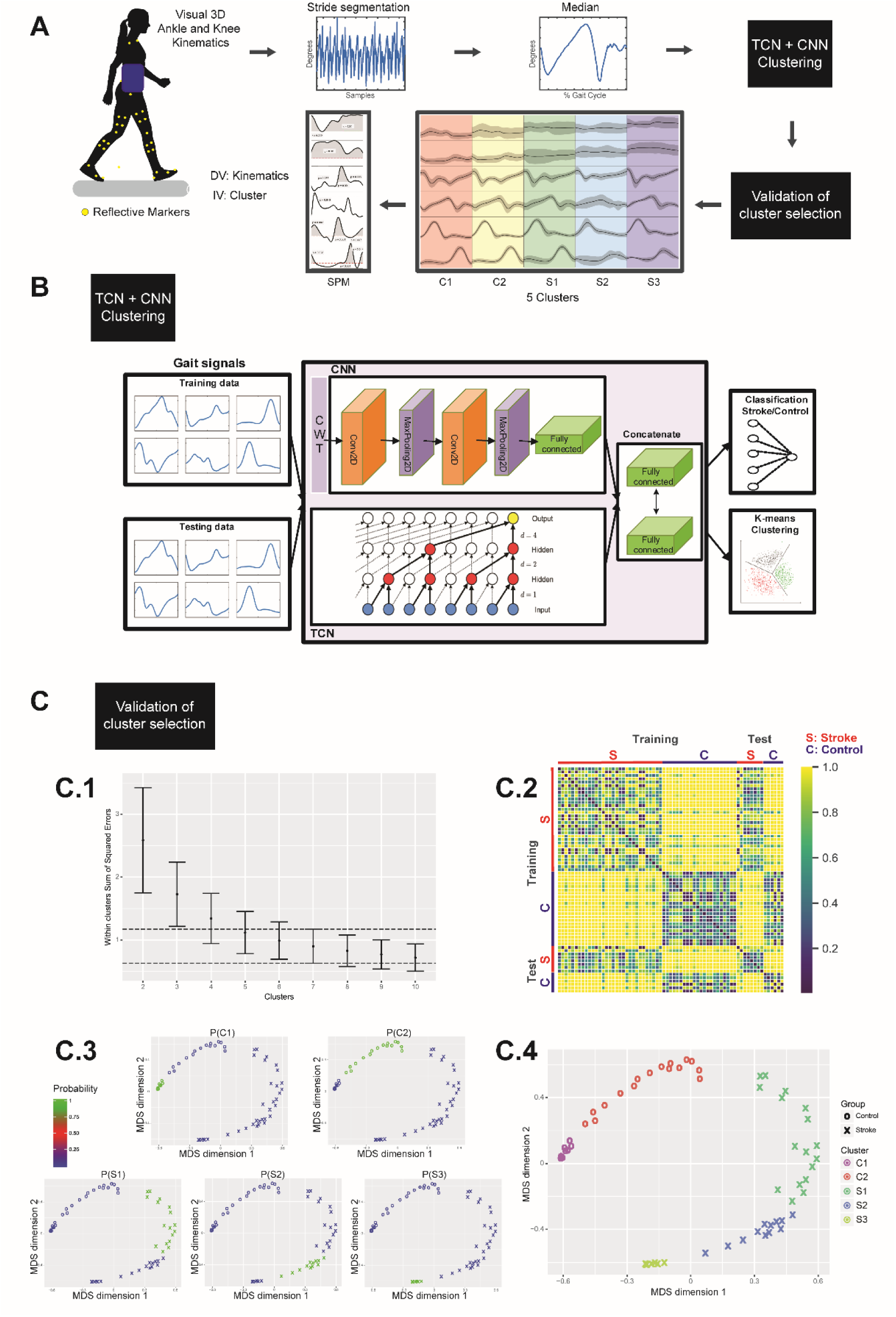
Study pipeline. **A**: Study pipeline. We used a lower extremity marker set to derive joint-level kinematics for the knee and ankle. We obtained an ensemble average of kinematics over the gait cycle, and this time series data was fed into a Convolution Neural Network (CNN)-Temporal Convolutional Network (TCN) ^54^, (Panel **B**) to obtain clusters of gait behaviors. We repeated the process 10000 times to assess the cluster number and composition (Panel **C**). We then used 1- D Statistical Parametric Mapping ^35^ to assess significant differences during the gait cycle between clusters for each kinematic variable. **B**: Detailed machine learning pipeline. CWT: Continuous Wavelet Transform. **C**: **C.1**: Optimal number of clusters using the one-standard error rule. **C.2**: Matrix of dissimilarities between each of the individuals, 1 indicates never in the same cluster, 0 always in the same cluster. **C.3**: Membership probability to each of the five clusters for every individual, in a multidimensional scaling 2D latent space (MDS dimension 1, MDS dimension 2) and a five-component Gaussian Mixture is computed. ^11^. **C.4**: A priori labeling in the MDS space. O: Control; X: Post-stroke; C1 and C2 are the control groups; S1, S2, and S3 are the post-stroke groups.

### Time series clustering analyses

#### Machine Learning Pipeline

We used a deep learning method combining a Convolutional Neural Network (CNN) ^28^ and a Temporal Convolutional Network (TCN) ^29^ to extract features from kinematic time series in the frequency and in the time domain, respectively, to have a complete gait representation for each individual. To extract the frequency-related features of the gait signals, we first applied a continuous wavelet transform ^30^ to express our data in the time-frequency domain and then used a CNN. In parallel, to extract the time-related features, we used a TCN on the kinematic time series. Hence, CNN and TCN were first trained with labels (control/stroke) to select features in a multivariate signal that can distinguish the two groups. Then, we used unsupervised time series k-means clustering to identify clusters of gait behaviors using the combined weights of the CNN-TCN features. The clustering pipeline was coded in Python (3.11, Python Software Foundation) and is available for download ^31^. In more detail, the different stages of our pipeline are (Fig. 1B):

1. Convolutional Neural Network ^32^: We first pre-processed the time series data into a continuous wavelet transform module that allows a two-dimensional representation of the signals as time-related frequency components. We multiplied each signal by the Morlet wavelet and the wavelet coefficients of the transformed signals were used as the inputs to the CNN. Then, the extracted wavelet coefficients are input into a CNN designed to learn spatial hierarchies of features automatically and adaptively through backpropagation. Two convolution kernels were applied to obtain 32 convolutional kernels with the size of 3×3, followed by a 2D max pooling layer, and 64 convolutional kernels with the size of 3×3, a 2D max pooling layer, are used to extract high-level features. A flattened layer was used to reorganize the feature maps into a one-dimensional array and fed into two consecutive fully connected dense layers of 128 and 64 features, respectively.
2. Temporal Convolutional Network ^29^: Since CNN only collects waveform characteristics, resulting in a lack of time series characteristics, the raw signals are fed in parallel, directly into a TCN module to extract temporal features. By fusing the two networks, we can effectively learn the spatial-temporal information in each gait cycle. The TCN is a residual network-based CNN designed for handling time sequence data. The output of TCN is a flattened layer of 64 features.
3. Supervised feature extraction: the outputs of CNN and TCN are concatenated into a combined TCN- CNN model and fed into two consecutive fully connected layers of 101 features and trained with the labels control/stroke to identify the high-level spatiotemporal features for neurotypical and post-stroke gait.
4. Unsupervised clustering: Once the full model is trained, it serves as a feature extractor for every participant. The gait signals are fed again into the trained model, then the extracted features are further used for time series k-means clustering.

To ensure that the results are representative of our dataset and not dependent on a unique training and testing split, we bootstrapped the classification stroke/control from CNN-TCN, CNN only and TCN only, a total of 10,000 times. At each iteration, a new random stratified 80/20 train/test split was performed to train the models. We also performed bootstrap analyses of the R time series clustering algorithms using 10,000 iterations, with each iteration being a new random seed to change the starting point of the algorithm.

### Number of clusters and cluster stability

To determine the optimal number of clusters c we calculated the within-clusters sum of squared errors over 10,000 bootstrap iterations ^33^ for 2 to 10 clusters and identified the number of clusters *c*^′^ that did not decrease the within-clusters sum of squared errors. We then selected *c* as the number of clusters having an average within-cluster sum of squared errors under one standard error of *c*^′ 11^ (Fig. 1 C.1)

After the bootstrap iterations, we obtained a clustering matrix (individuals x iterations) where each element represents a cluster number from 1 to . To assess the occurrences of individuals classified together in the same cluster (Fig. 1 C.2), we first calculated a similarity matrix (individuals x individuals):

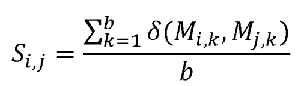

Where and are indices to represent individuals, is the iteration index and

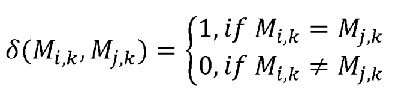

Then to recover the number of stable clusters, we compute the dissimilarity matrix *D* as *D*=1−*S*, and projected it in a lower two-dimensional latent space using multidimensional scaling, preserving the pairwise Euclidean distance between each element ^11^ (Fig. 1 C.3).

Finally, with the new latent space coordinates, we performed a clustering for clusters using a Gaussian mixture model as a sum of Gaussian distributions with their own mixing proportions, mean, and full covariance matrices ^11^. This probabilistic method is better suited to handle nonspherical clusters as we obtained in the latent space (Fig. 1 C.3) and allows us to obtain individual probability estimates of each participant belonging to each of the identified clusters ^11^.

The Gaussian mixture clustering was performed times, each iteration corresponding to new initialization parameters for the Gaussian mixture model. The final clustering model was chosen to minimize the Akaike Information Criterion across the iterations (Fig. 1 C.4) ^11,34^.

### Pipeline comparison

We verified if combining both frequency and time-related features improves the classification into stroke/control by comparing our full CNN-TCN pipeline to the classification output of CNN only and TCN only. At the supervised stage, i.e., stage 3, after the last layer, we added a size 1 fully connected layer using a sigmoid activation for a binary stroke/control classification. We processed similarly to assess the classification of the CNN and TCN blocks alone. We used the classification of each test set to build the confusion matrix of actual versus predicted labels (stroke/control) and compute a standard accuracy metric of the supervised models ^28^. We also compared our method to a readily available unsupervised multivariate time series clustering using a dynamic time-warping algorithm (R package *dtwclust*) ^13^ and a partitional clustering performed on the dynamic time-warping barycenter averaging centroids. We used the entire set for the unsupervised clustering into two groups.

Finaly, we evaluated the goodness of fit of the five components Gaussian mixture model by comparing the Akaike Information Criterion for our CNN-TCN pipeline compared to TCN only and CNN only. In addition, we also verified that the supervised learning layers are not biased toward classifying an individual into a control or a stroke group (Supplementary materials 1). To do so, we looked at the projection of the dissimilarity matrix into the multidimensional scaling space and after the 10,000 iterations when (1) we mislabeled a stroke participant that had the highest Fugl-Meyer score (i.e. 34) into a control participant (total 67 individuals, 38 post-stroke and 29 control), and (2) we added the high Fugl-Meyer stroke participant with a control and a stroke label (total 68 individuals, 39 post-stroke and 29 control).

### Statistical analysis

#### Comparisons between individuals post-stroke and neurotypical controls

We assessed for significant differences in demographics and clinical measures between participants post-stroke and neurotypical controls in SPSS (29.0, IBM Corp, Armonk, USA). Data were assessed for normality using the Shapiro-Wilk test. If data were normally distributed, we used independent samples t-tests to compare data between groups. Otherwise, we used the Wilcoxon Signed Rank test. To compare differences in the distribution of males vs. females across groups, we used a Chi-Square test. For normal data, values are reported as mean ± SD, and for non-normal data, values are reported as median ± IQR.

#### Comparison within clusters

Within each control cluster (C1 and C2), we compared the participants’ self-selected speed to the matched walking speed using a paired t-test.

Within each stroke cluster (S1, S2, S3), we used Student’s t-tests to assess for asymmetries within clusters between paretic/non-paretic extremities for step length, swing time, and stance time. The significance level was Bonferroni corrected according to the number of tests. We also computed Pearson’s correlation coefficient between the treadmill walking speed and the clinical assessments.

#### Comparison between clusters

We used a one-way ANOVA to compare demographics, clinical measures, and spatiotemporal gait measures (dependent variables) between clusters (independent variable). If we observed significant results from the ANOVA, we performed multiple comparisons with Tukey’s test. We

clarify that these discrete metrics were not used in the machine learning pipeline to identify the distinct gait clusters and were assessed post-hoc.

We used 1-dimensional statistical parametric mapping ^35^ one-way ANOVA with cluster identifier as the independent variable to assess kinematic time series differences between clusters. Post-hoc tests were done via statistical parametric mapping t-tests with Bonferroni correction for multiple comparisons. We used 2-dimensional statistical parametric mapping ^36^ to compare the continuous wavelet transform matrix coefficients between groups in the time-frequency domain (Supplementary materials 2). These kinematics characteristics were used by the machine learning pipeline to identify the distinct gait clusters.

## Results

Participants post-stroke were 59.5 ± 10.8 years old, had a mass of 76.5 ± 15.8 kg, and were 92 ± 84.5 months post-stroke. 22 participants were male and 17 female. Control participants were 62.4 ± 14.2 years old and had a mass of 74.1 ± 16.3 kg. 12 control participants were male and 16 female (Table 1). As data were extracted for the middle 50 s of walking to ensure a steady gait pattern, this resulted in 39.2 ± 7.2 strides for control participants at the matched speed and 37.9 ± 6.5 strides for individuals post-stroke. Two participants post-stroke wore an ankle brace while walking.

### No significant differences in demographics between participants post-stroke and neurotypical controls

We observed no significant differences between participants post-stroke and neurotypical controls in age, height, mass, self-selected treadmill walking speed, or matched walking speed (p>0.050). We did not observe differences in the study samples’ proportion of males vs. females (p>0.05). We observed significant differences in ABC and BBS between participants post-stroke and neurotypical controls (Wilcoxon Signed Rank test p<0.001, Table 1).

### CNN-TCN performed better than other time series clustering algorithms

The confusion matrix for the four evaluated algorithms is presented in Table 2. During the supervised analyses and over 10000 iterations, our CNN-TCN pipeline predicted individuals of the test set correctly 85.0 ± 14.7 % of the time for stroke participants and 87.7 ± 11.2 % for control participants. The overall accuracy for the CNN-TCN model was 86.4%, slightly higher than for CNN alone (86.3%) and TCN alone (84.1%) to classify individuals as stroke/control.

**Table 2:** Confusion matrix. Confusion matrix for individual blocks of Convolutional Neural Network (CNN), Temporal Convolutional Network (TCN), dual-stage CNN-TCN and Dynamic Time Warping clustering (dtwclust)

| CNN |  | Predicted |  |
| --- | --- | --- | --- |
|  |  | Stroke | Control |
| Actual | Stroke | 85.4% | 14.5% |
|  | Control | 12.9% | 87.1% |

| TCN |  | Predicted |  |
| --- | --- | --- | --- |
|  |  | Stroke | Control |
| Actual | Stroke | 83.0% | 17.0% |
|  | Control | 14.8% | 85.2% |

| CNN-TCN |  | Predicted |  |
| --- | --- | --- | --- |
|  |  | Stroke | Control |
| Actual | Stroke | 85.0% | 15.0% |
|  | Control | 12.3% | 87.7% |

| dtwclust |  | Predicted |  |
| --- | --- | --- | --- |
|  |  | Stroke | Control |
| Actual | Stroke | 52.5% | 48.5% |
|  | Control | 5.2% | 94.8% |

However, the Akaike Information Criterion was lower for the CNN-TCN model (-142.5) compared to CNN (6.8) alone and TCN alone (-64.0), indicating a significantly better fit of the five component Gaussian Mixture Model for our proposed full CNN-TCN pipeline.

The unsupervised clustering based on dynamic time warping package in R was better at predicting control individuals (94.8 ± 0.1 %) but was close to random at predicting stroke individuals (52.5 ± 0.1 %), which resulted in worse overall accuracy (73.7%). Thus, our fused pipeline performed better than the individual components of the pipeline and the R package during supervised analyses using two classes due to the added ground truth knowledge and the combination of both temporal and spatial features.

### k=5 provided stable clusters

When determining the optimal number of clusters, we first identified the “elbow” ^33^ at *K*^′^=7, and then selected *K*=5 as the optimum number of clusters using the one standard error (Fig. 1 C.1). Thus, we computed a 5-component Gaussian mixture model from the dissimilarity matrix (Fig. 1 C.2). We observed two control clusters (C1, C2) and three clusters of individuals post-stroke (S1, S2, S3) (Fig. 1 C.3 and C.4). The mixing proportion, which indicates cluster membership likelihood, of almost all individuals but one to be assigned to their final cluster was above 99% (Fig. 1 C.3). Only one participant in S2 had a mixing proportion of 93% for S2, with a mixing proportion of 7% for S1. None of the individuals post-stroke were assigned to a control cluster (Fig. 1 C.4).

We further assessed that our pipeline did not introduce bias in the supervised stage with the added knowledge of groups to extract features. The participant with a maximal Fugl-Meyer, thus considered to have recovered the most motor function among our participant ^21^, was always projected in the stroke clusters part of the multidimensional scaling space when mislabeled to control or added with a stroke and control label (Supplementary materials 1).

### All clusters had significantly different clinical characteristics and kinematic patterns

We compared demographics, clinical measures, and spatiotemporal characteristics using one-way ANOVA between the five identified clusters (C1, C2, S1, S2, S3), except for FM and FGA which was between the post-stroke clusters only (S1, S2, S3). All clinical measures were significantly different between groups: FM (p=0.003), FGA (p=0.02), ABC (p<0.001), BBS (p<0.001). Age (p=0.56) and height (p=0.94) were not different between clusters. The following spatiotemporal characteristics were significantly different between clusters (Fig. 3): walking speed (p=0.003), stride length (p=0.008), paretic/non-dominant step length (p=0.010), non-paretic/dominant step length (p=0.007), stride time (p=0.001), paretic/non-dominant stance time (p<0.001), non-paretic/dominant stance time (p=0.004). Paretic/non-dominant swing time (p=0.57) and non-paretic/dominant swing time (p=0.08) were not significantly different between clusters.

For the knee and ankle kinematics, using 1D SPM one-way ANOVA, we observed significant differences between clusters in all degrees of freedom bilaterally: paretic/non-dominant ankle plantar/dorsiflexion (p<0.001 whole gait cycle), non-paretic/dominant ankle plantar/dorsiflexion (p=0.049 at loading response, p=0.001 during pre-swing and p=0.006 at terminal stance), paretic/non-dominant ankle abduction/adduction (p=0.03 during loading response), non-paretic/dominant ankle abduction/adduction (p<0.001 whole gait cycle), paretic/non-dominant knee flexion/extension (p=0.03 during initial contact and loading response, p<0.001 during pre-swing, and p=0.04 during terminal swing) and non-paretic/dominant knee flexion/extension (p=0.002 during terminal swing, p=0.001 during loading response, and swing p=0.007 during pre-swing). We describe each cluster next:

**Control Cluster 1 (C1):** N=12. Normative gait pattern (Fig. 2). Participants in C1 were 62.8 ± 13.8 years old, walked at a self-selected speed of 0.96 ± 0.28 m/s, and were significantly slower when matched to participants post-stroke (0.65 ± 0.20 m/s, p=0.010). This cluster was composed exclusively of control individuals. Their non-dominant and dominant step lengths were both 0.39 ± 0.07 m, stride length was 0.78 ± 0.13 m, stride time was 1.45 ± 0.26 s, non-dominant stance time was 0.99 ± 0.22 s, dominant stance time was 1.03 ± 0.26 s, non-dominant swing time was 0.46 ± 0.06 s, and dominant swing time was 0.45 ± 0.06 s (Fig. 3). The kinematics in this cluster are those described in the literature for non-injured, neurotypical adults ^37–40^. We report all kinematic post-hoc comparisons relative to this cluster (blue line in Fig. 4 in all panels).

**Figure 2:**
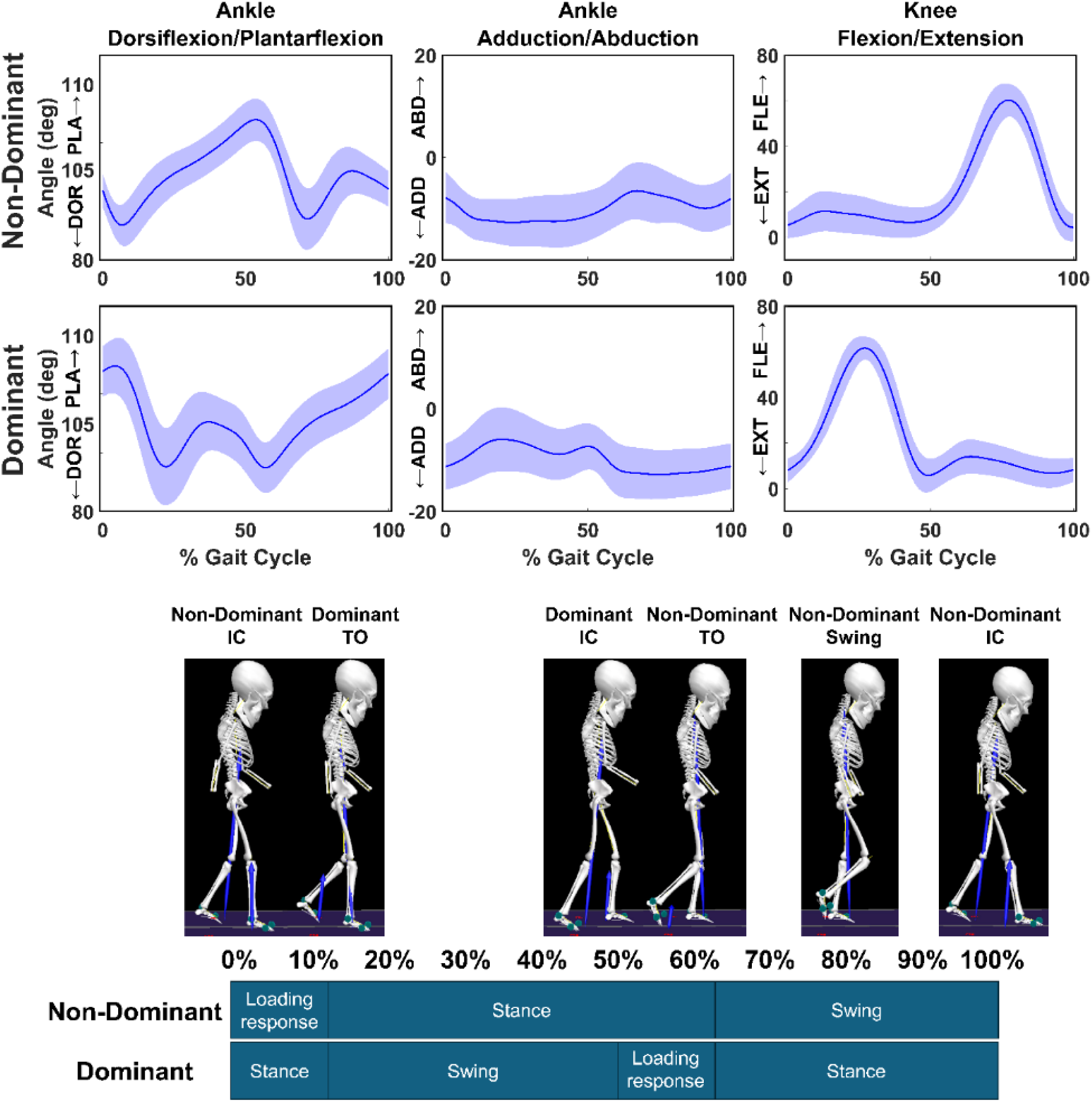
Normative gait cluster (C1). Gait kinematics of the normative walking cluster (C1) and corresponding gait phases snapshots from Visual 3D. The cycle starts at the non-dominant initial contact (IC). TO: toe-off.

**Figure 3:**
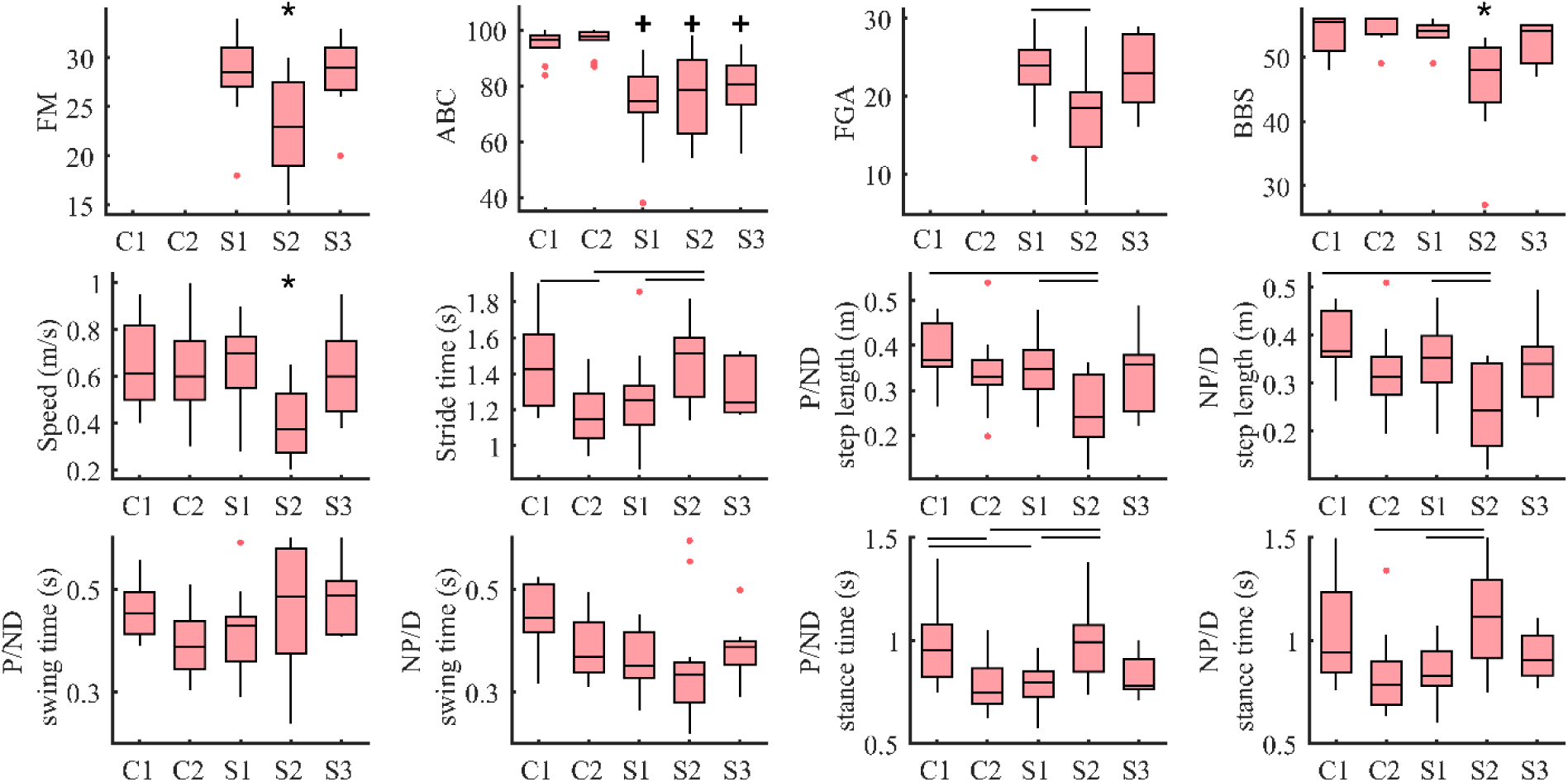
Spatiotemporal and clinical measures between clusters. Clinical assessments and gait spatiotemporal characteristics between the five clusters (Controls: C1 and C2, Stroke: S1, S2, and S3). The non-dominant side for control individuals is compared to the paretic side of post-stroke individuals, and dominant to paretic. One-way ANOVA showed a difference for all parameters (p<0.05), except paretic/non-dominant and non-paretic/dominant swing times (p=0.574 and p=0.078 respectively). Tukey’s test was used as post-hoc. FM: Fugl-Meyer lower extremity, ABC: Activities Balance Confidence, FGA: Functional Gait Assessment, BBS: Berg Balance Scale, P: Paretic, NP: Non-Paretic, D: Dominant, ND: Non-Dominant. * lower than all the other clusters, + lower than C1 and C2, ⎯ difference between two clusters. The significance level is set at p<0.05.

**Figure 4:**
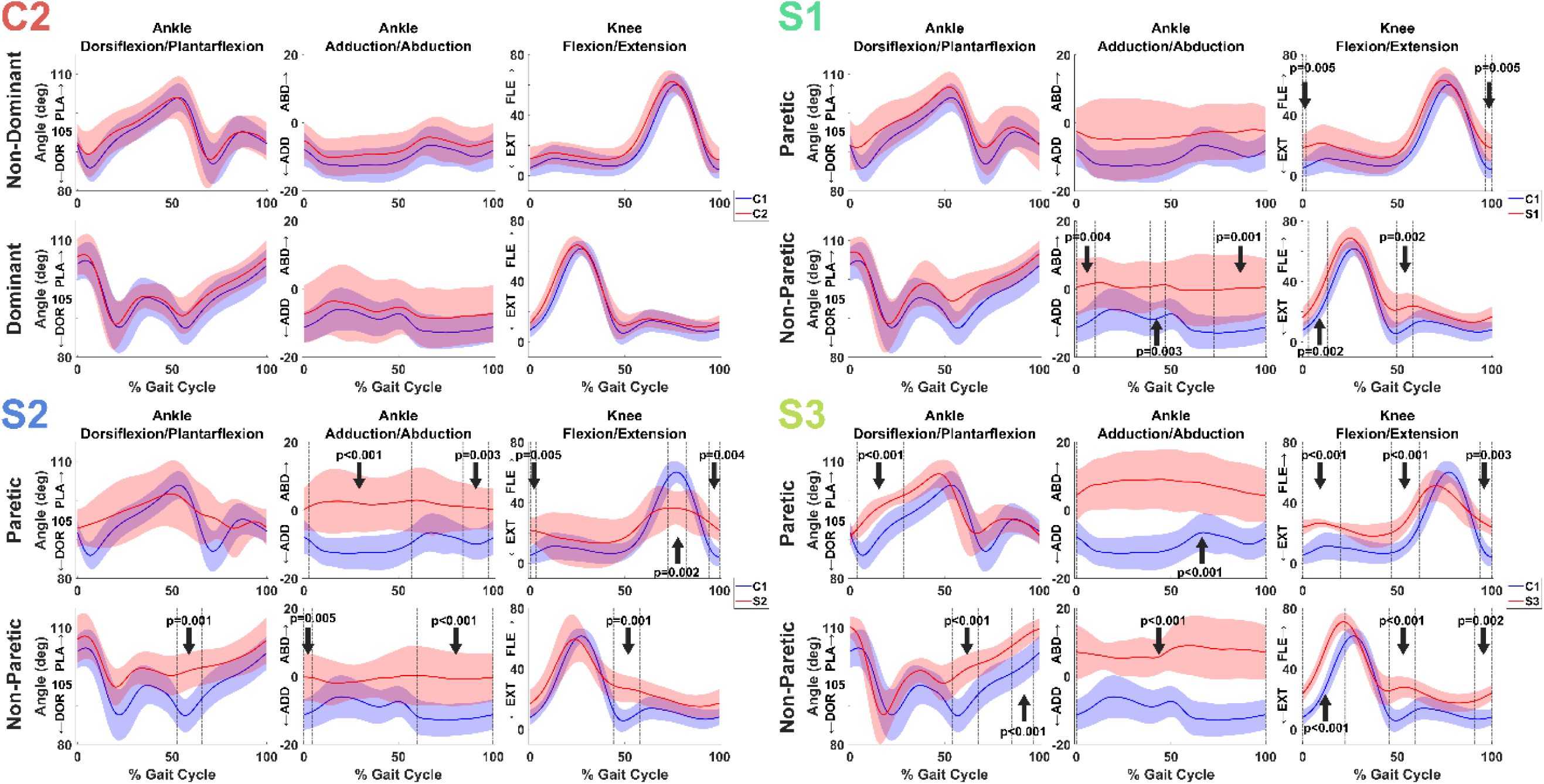
1D Statistical Parametric Mapping. Post-hoc 1D-Statistical Parametric Mapping t-test with Bonferroni correction of joint kinematics for the median gait cycle between the reference control cluster in blue (C1) and the other clusters in red (Controls: C2, Stroke: S1, S2, and S3). The non- dominant side for control individuals is compared to the paretic side of post-stroke individuals, and dominant to paretic. The vertical dashed lines surrounding a vertical arrow indicate endpoints of significant differences during the gait cycle.

**Control Cluster 2 (C2):** N=16. Control participants with short stride times. This cluster was also composed exclusively of control individuals. Participants in C2 were 62.1 ± 15.6 years old, walked at a self-selected speed of 0.75 ± 0.22 m/s (significantly slower than the self-selected speed of C1, p=0.03), and a matched speed to participants post-stroke of 0.63 ± 0.18 m/s, (significantly slower than their self-selected speed, p=0.034, but not significantly slower than the imposed speed of C1). Compared to C1, they had shorter non-dominant stance time (0.78 ± 0.13 s, p=0.02) and stride time (1.18 ± 0.16 s, p=0.009) when walking at speeds matched to participants post-stroke (Fig. 3). The non-dominant step length (0.34 ± 0.08 m), dominant step length (0.32 ± 0.08 m), stride length (0.66 ± 0.16 m), dominant stance time (0.82 ± 0.18 s), non-dominant swing time (0.39 ± 0.06 s), and dominant swing time (0.35 ± 0.15 s) were not significantly different from C1. Interestingly, we did not observe significant differences in kinematics between C1 and C2 (Fig. 4). The differences between the two control clusters are present in the time-frequency domain for the dominant knee flexion/extension (Supplementary materials 2). In the same range of normalized frequency [0.025-0.034 cycles/sample], compared to C1, the coefficients of the continuous wavelet transform matrix in C2 were lower during initial swing and loading response, but higher during mid swing (Supplementary materials 2).

**Stroke Cluster 1 (S1):** N=17. Stroke participants with increased knee flexion at initial contact/loading response bilaterally. Participants in S1 were 58.2 ± 13.8 years old and walked at a speed of 0.65 ± 0.17 m/s (Fig. 3). Their FM score was 29 ± 4, indicating mild impairment, FGA was 23 ± 5, and BBS was 54 ± 2. ABC was 74 ± 14, lower than C1 (p<0.001) and C2 (p<0.001) (Fig. 3). Compared to C1, they had a shorter paretic stance time (0.76 ± 0.19 s, p=0.005), but no differences were found for paretic step length (0.35 ± 0.07 m), non-paretic step length (0.35 ± 0.08 m), stride length (0.70 ± 0.15 m), stride time (1.24 ± 0.22 s), non-paretic stance time (0.88 ± 0.21 s), paretic swing time (0.48 ± 0.30 s), non-paretic swing time (0.37 ± 0.06 s) (Fig. 3). No asymmetry was detected between the paretic and non-paretic extremities for step length (p=0.90), swing time (p=0.14) and stance time (p=0.07). Compared to C1, we observed increased paretic knee flexion at initial contact/loading response (p=0.005), and during terminal swing (p=0.005) (Fig. 4). In the non-paretic extremity, we observed increased ankle abduction during pre-swing (p=0.004), terminal swing (p=0.003), and terminal stance (p=0.001). Finally, for the non-paretic side, we observed increased knee flexion during pre-swing (p=0.002), at initial contact, and loading response (p=0.002) (Fig. 4). This cluster had a significant positive correlation between walking speed and FM (r=0.77, p<0.05), walking speed and FGA (r=0.69, p=0.003), but no correlation between speed and ABC or BBS (Fig. 5). One participant in S1 wore an ankle brace.

**Figure 5:**
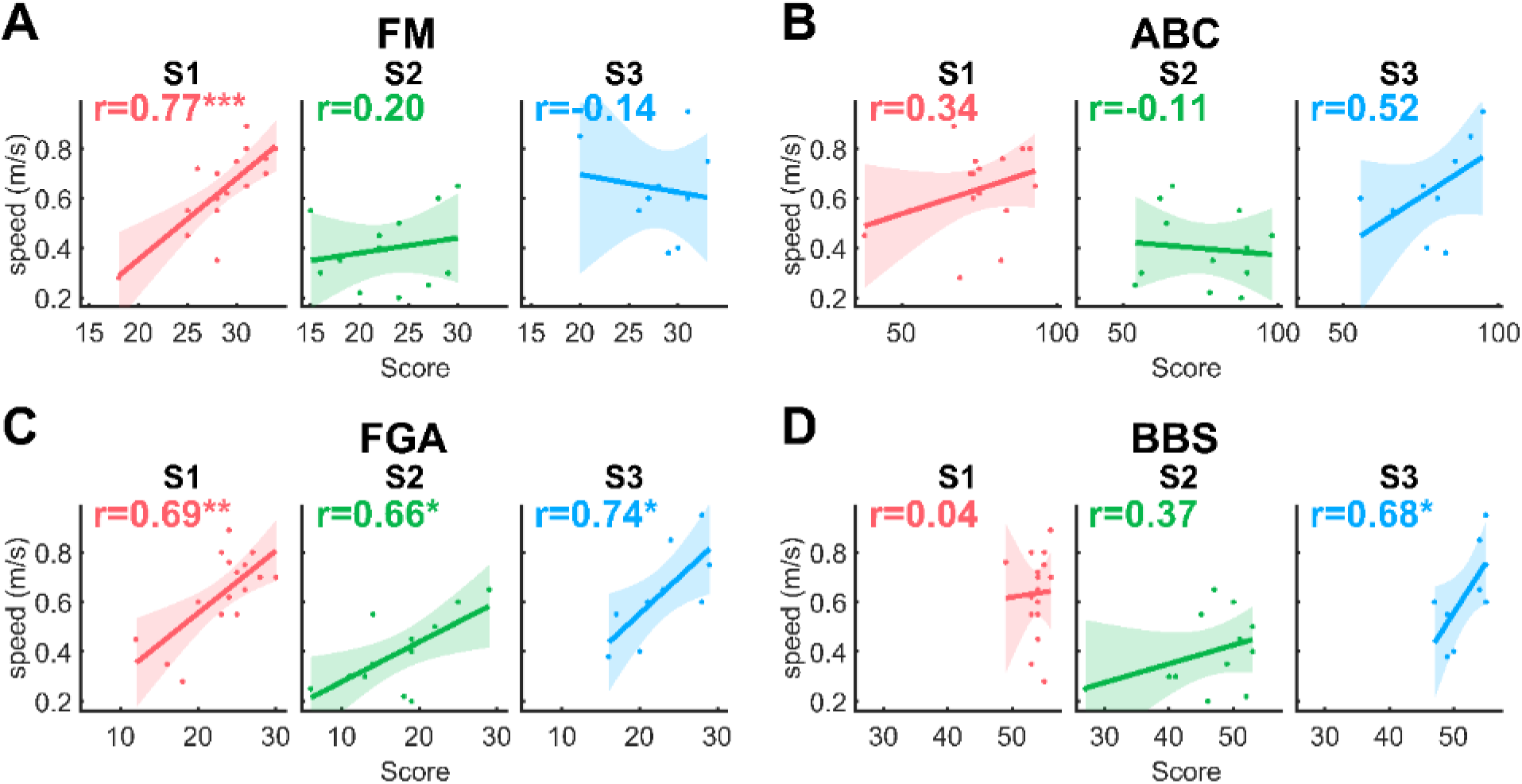
Correlations. Correlations between walking speed and clinical assessment scores within post-stroke clusters (S1 in red, S2 in green, S3 in blue). FM: Fugl Meyer lower extremity, ABC: Activities Balance Confidence, FGA: Functional Gait Assessment, BBS: Berg Balance Scale. The significance level is set at p<0.05. ***: p<0.001, **: p<0.01, *: p<0.05

**Stroke Cluster 2 (S2):** N=12. Increased stance knee flexion bilaterally and reduced paretic swing knee flexion. Participants in S2 were 64.0 ± 3.4 years old and walked at a slower speed (0.40 ± 0.15 m/s, p<0.05 compared to all other clusters) (Fig. 3). The FM score was 23 ± 5 indicating moderate impairment. FM score was lower than S1 (p=0.004) and S3 (p=0.02). FGA was 17 ± 6, lower than S1 (p=0.019) but not S3 (p=0.06). BBS was 46 ± 7, lower than S1 (p<0.001) and S3 (p=0.02). ABC was 76 ± 15. Compared to C1, individuals post-stroke in S2 had a shorter paretic step length (0.26 ± 0.08 m, p=0.005), non-paretic step length (0.25 ± 0.09 m, p=0.004), and stride length (0.51 ± 0.17 m, p=0.004). Compared to S1, paretic step length (p=0.005), non-paretic step length (p=0.004) and stride length (p=0.004) were also shorter, with longer stride time (1.47 ± 0.22 s, p=0.039) and paretic stance time (0.99 ± 0.18 s, p=0.005).

Their non-paretic stance time was 1.12 ± 0.25 s, paretic swing time 0.48 ± 0.15 s, non-paretic swing time 0.35 ± 0.12 s (Fig. 3). No asymmetry was detected between the paretic and non-paretic extremities for step length (p=0.84), swing time (p=0.07) and stance time (p=0.18).

Compared to C1, we observed increased paretic ankle abduction during loading response and stance (p<0.001), and terminal swing (p=0.003). Compared to C1, we observed increased paretic knee flexion at initial contact and loading response (p=0.005), and during terminal swing (p=0.004), and decreased paretic knee flexion mid-swing (p=0.004) (Fig. 4). In the non-paretic extremity, we observed increased ankle abduction during the entire stance phase (p<0.001) and pre-swing (p=0.005) (Fig. 4). We also observed decreased non-paretic dorsiflexion at initial contact and loading response (p=0.001). Finally, we observed increased non-paretic knee flexion from terminal swing to loading response (p=0.001) (Fig. 4). This cluster had a significant positive correlation between walking speed and FGA (r=0.66, p=0.02) only (Fig. 5), potentially indicative of balance impairment.

**Stroke Cluster 3 (S3):** N=10. Increased ankle abduction. Participants in S3 were 56.7 ± 11.1 years old and walked at a speed of 0.62 ± 0.19 m/s. The FM score was 28 ± 4, ABC was 79 ± 12, FGA was 23 ± 5, BBS was 52 ± 3 (Fig. 4). Compared to C1, they had similar gait spatiotemporal characteristics paretic step length was 0.33 ± 0.09 m, non-paretic step length 0.34 ± 0.08 m, stride length 0.67 ± 0.17 m, stride time 1.31 ± 0.22 s, paretic stance time 0.82 ± 0.18 s, non-paretic stance time 0.92 ± 0.12 s, paretic swing time 0.49 ± 0.07 s, non-paretic swing time 0.38 ± 0.05 s. The peak knee flexion occurring earlier in the gait cycle caused a longer swing time on the paretic side (p=0.001). Despite no significant differences in clinical scores compared to S1 (Fig. 3), this cluster showed different kinematic impairments (Fig. 3). Compared to C1, we observed increased paretic ankle abduction bilaterally for the entire gait cycle (both p<0.001) (Fig. 4). We observed decreased paretic dorsiflexion during loading response and mid-stance (p<0.001) (Fig. 4). We observed increased paretic knee flexion from loading response to mid-stance (p<0.001) and during pre-swing (p<0.001) and terminal swing (p=0.003), Fig. 4. In the non-paretic extremity, we observed decreased dorsiflexion during loading response (p=0.001) and terminal stance (p=0.001) (Fig. 4). Finally, we observed increased knee flexion during pre-swing and initial swing (p<0.001), initial contact and loading response (p<0.001), and terminal stance (p=0.002) (Fig. 4). This cluster had a significant positive correlation between walking speed and FGA (r=0.66, p=0.02), walking speed and BBS (r=0.68, p=0.04), but no correlation between speed and FM or ABC (Fig. 5). One participant in S3 wore an ankle brace.

## Discussion

Gait impairment is heterogeneous, posing a challenge in the prescription of research or rehabilitation interventions ^41^. To inform rehabilitation, previous research identified subgroups of gait behaviors post-stroke based on peak spatiotemporal characteristics, peak kinematics, peak kinetics, or muscular activity ^1,2,5,6,8^. These discrete metrics cannot capture the simultaneous temporal and spatial variation in the gait cycle seen in people post-stroke. Here, we developed a pipeline using convolutional networks to identify subgroups of gait behaviors and tested it with kinematic time series data of neurotypical and chronic stroke individuals. We showed that providing the true labels in a supervised stage first to extract frequency and time-related gait features from kinematic time series was more advantageous than fully unsupervised time series clustering techniques.

Our pipeline identified distinct walking behaviors both in neurotypical control and post-stroke participants. In neurotypical participants, the subgroups were differentiated by their self-selected walking speed which was slower in C2, thus shaping their walking pattern by altering spatiotemporal and kinematics characteristics ^42^. In participants post-stroke, the subgroups were characterized by kinematic impairments that differentially affected distinct phases of the gait cycle. Contrary to our previous work that used only peak kinetics and spatiotemporal characteristics ^5^, our results indicate that at a joint kinematics level, post-stroke participants are distinct from neurotypical controls, even when they have minimal impairment measured via clinical outcomes. Our results also indicate that individuals post-stroke can show similar levels of function and impairment measured using clinical outcomes while displaying vastly different joint kinematics. Finally, our results also highlight movement patterns in the non-paretic extremity function during gait which are often overlooked, seldom reported ^6,43,44^ and that differ from the typically described compensatory pattern ^45^. Using our pipeline, we have provided a more detailed assessment of the distinct types of gait behaviors post-stroke, which affect both the paretic and non-paretic extremities, and can point at intervention targets post-stroke.

The three post-stroke clusters obtained in our study point to different impairments and potential rehabilitation interventions. S1, stroke participants with increased knee flexion at initial contact/loading and terminal swing, response bilaterally showed the least amount of gait impairments. The increased paretic knee flexion seen during loading response and terminal swing corresponds to common knee patterns post-stroke ^6,46^ and might indicate potential hip extensor weakness ^43,44^. Treatment for participants in S1 might include functional step training. Participants in S1 had similar clinical and spatiotemporal characteristics as ‘*the moderate speed, symmetric, and short stance times’* cluster in our previous work ^5^. S2, post-stroke participants with increased stance knee flexion, reduced swing knee flexion, and reduced dorsiflexion showed the most impaired gait pattern. Participants in this group also showed increased ankle abduction in the paretic extremity, which might point to limb circumduction to advance the paretic limb forward due to the observed decreased knee flexion during the swing phase ^6,44,46^. This group’s spatiotemporal characteristics are quasi-similar to the *‘slow speed and frontal plane force asymmetries’* group in our previous study, with only differing stance time asymmetry not present in S2. Participants in this group would benefit from dorsiflexion strengthening, electrical stimulation of dorsiflexors and during swing to elicit a mass flexion response ^41,44^, ankle-foot orthosis ^41,44^, manual cues to guide knee flexion during swing ^41,47^, and balance training ^48^. S3 showed no differences in speed, FM, FGA, or Berg to S1, yet it showed additional gait impairments, including a flexed knee during stance bilaterally, increased non-paretic knee flexion during swing, reduced dorsiflexion bilaterally, and increased ankle abduction. Potential treatments for participants in S3 include strengthening of hip extensors and hamstrings, gait retraining with an emphasis on improving motor control and coordination.

Previous studies have used speed alone ^3^, or identified speed as the main determinant of cluster allocation ^2^. Our findings contrast these findings, as S1 and S3 had similar average speeds and clinical characteristics while showing different kinematic patterns indicating that clinical scores are not granular enough to show specific gait kinematic impairments. This was also confirmed by the heterogeneity of correlation between speed and clinical scores within the clusters. Using time series joint kinematics, we obtained two moderate-speed clusters and one slow cluster. Some of the characteristics of our clusters resembled those previously reported ^2^. For example, S1, which had slightly decreased knee extension in terminal swing, initial contact and loading response but adequate dorsiflexion, resembled the Fast group reported by Mulroy ^2^, despite the more moderate walking speed in our participants. S2 had a slow velocity with excessive knee flexion in stance and inadequate dorsiflexion in swing, similar to what was reported by Mulroy ^2^ as the slow flexed group ^2^. We supplement this information by showing impaired non-paretic kinematics, particularly increasing non-paretic ankle abduction through stance, and reduced non-paretic dorsiflexion during swing, and increased non-paretic knee flexion in loading response in this group. We did not observe a knee hyperextension pattern as in previous work ^2,6,46^. Overall, our findings show that clinical measures such as speed or FM score are not sensitive to kinematic differences, and thus our approach can provide additional insights beyond what is provided by clinical measures.

Combining both time-frequency (CNN) and time-related features (TCN) to find subgroups of gait did not improve accuracy to classify neurotypical and post-stroke gait but provided a better model to identify clusters of gait behaviors post-stroke. While kinematics differences were expected in the post-stroke clusters ^2,5,6^, they were not significant in the two control clusters. By adding the time-frequency domain in our pipeline, we were able to differentiate 1) between neurotypical adults presenting normative gait patterns and neurotypical adults with affected kinematics because of walking at a matched slower speed ^49^, 2) between highly functional post-stroke individuals who scored high in all the clinical assessments and ‘visually normal’ kinematics and neurotypical controls. Our approach has the potential to complement and augment the clinical observation of gait since it can use any type of time series captured in a clinical setting with wearable devices ^50^ or phones ^51^.

### Limitations

The greatest limitation of our study is the lack of hip kinematics used in the definition of gait clusters. The lack of these measures limits our ability to identify subtypes of hip gait impairment blurring our understanding of the causes of knee and ankle impairment, which might originate due to impaired hip function. The inclusion of hip kinematics might lead to detecting additional clusters, and further expand the implications of our work, such that our current and future work will require the inclusion of hip kinematics. Participants were allowed to use walking aids, which modify ankle and knee kinematics post-stroke ^52^. Given that only two participants in our study wore ankle braces, this should not affect our findings significantly. The spasticity of participants post-stroke was not controlled for, and no information was collected regarding botulinum toxin injection to treat lower limb spasticity, which may also affect gait ^53^. Another limitation of our work is the use of a median gait cycle for each participant. Initial attempts were made to use non-segmented time series data collected for 30 seconds, but we ran into issues with autoencoder clustering as no two participants were alike using this approach, and even the supervised initial analyses using a single stroke and a single control cluster performed with an accuracy below chance. A final limitation is that we only measured self-reported injuries in control participants, and thus, the presence of two control clusters might be due to underlying injuries or impairment in control individuals, which may not be fully accounted for in our demographics.

## Conclusion

The presented machine-learning pipeline used kinematic time series to identify five distinct subgroups of gait behavior. We showed that individuals post-stroke were clearly different from neurotypical individuals at a joint level, even when they had mild impairment and similar spatiotemporal characteristics. Our approach has the potential to aid clinicians by augmenting observation of gait. Moreover, it can be applied to any type of pathology affecting gait and any type of one-dimensional data collected during gait.

## List of abbreviations

EMG: electromyography
BBS: Berg Balance Scale (max 56)
ABC: Activities Balance Confidence score (max 100)
FM: Fugl-Meyer Assessment (Lower extremities, max 34)
FGA: Functional Gait assessment (max 30)
CNN: Convolutional Neural Network
TCN: Temporal Convolutional Network
C1: Control cluster 1
C2: Control cluster 2
S1: Stroke cluster 1
S2: Stroke cluster 2
S3: Stroke cluster 3

## Declarations

### Ethics approval and consent to participate

Data collection was approved by the University of Southern California Institutional Review Board, with numbers HS-19-00430 and HS-18-00533, and each participant provided written informed consent. The University of Southern California Institutional Review Board IRB HS-19- 00075 approved data curation across multiple studies. The Chapman University IRB-23-12 approved de-identified data transfer from USC to Chapman University and secondary data analyses. All study aspects conformed to the principles described in the Declaration of Helsinki.

### Availability of data and materials

The data and machine learning code in Python for the current study are available for download in a public repository ^31^. A short video (Supplementary Material 3) is also provided to illustrate gait differences between the clusters ^31^.

## Funding

This work was supported by grants NIH NCMRR R03HD107630 and NIH NCATS R03TR004248 and KL2TR001854 to N. Sánchez (NSa).

### Author contributions

AK: Conceptualization, Methodology, Software, Formal analysis, Writing – Original Draft, Visualization. AM: Conceptualization, Writing – Review & Editing. NSc: Conceptualization, Methodology, Writing – Review & Editing. JM: Funding acquisition, Resources, Writing – Review & Editing. YW: Conceptualization, Methodology, Software, Visualization, Writing – Original Draft. NSa.: Funding acquisition, Resources, Supervision, Investigation, Conceptualization, Visualization, Formal analysis, Writing – Original Draft, Data Curation

## Acknowledgments

We want to thank Chang Liu, PhD, Sungwoo Park, PhD, Tara Cornwell, Ryan Novotny, Catherine Yunis, and Isabel Munoz-Orozco who contributed to data collection and processing for this study.

---

**Supplementary materials 1.**
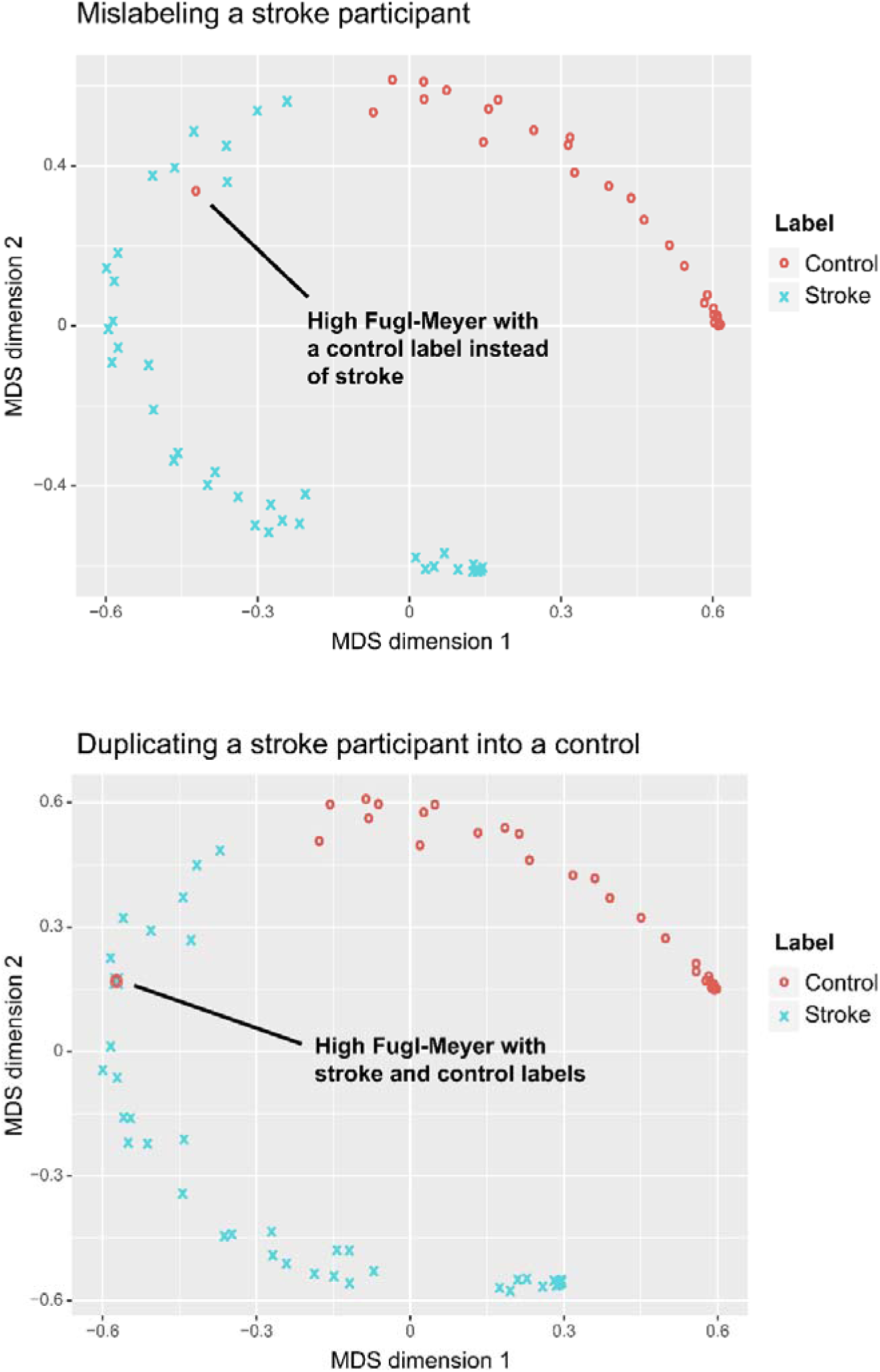
Projection of each individual into the multidimensional scaling latent space. When the highest Fugl-Meyer score participant is mislabeled as a control and when we input this participant with a stroke and a control label, our pipeline still projects the participant next to other people post-stroke.

**Supplementary materials 2.**
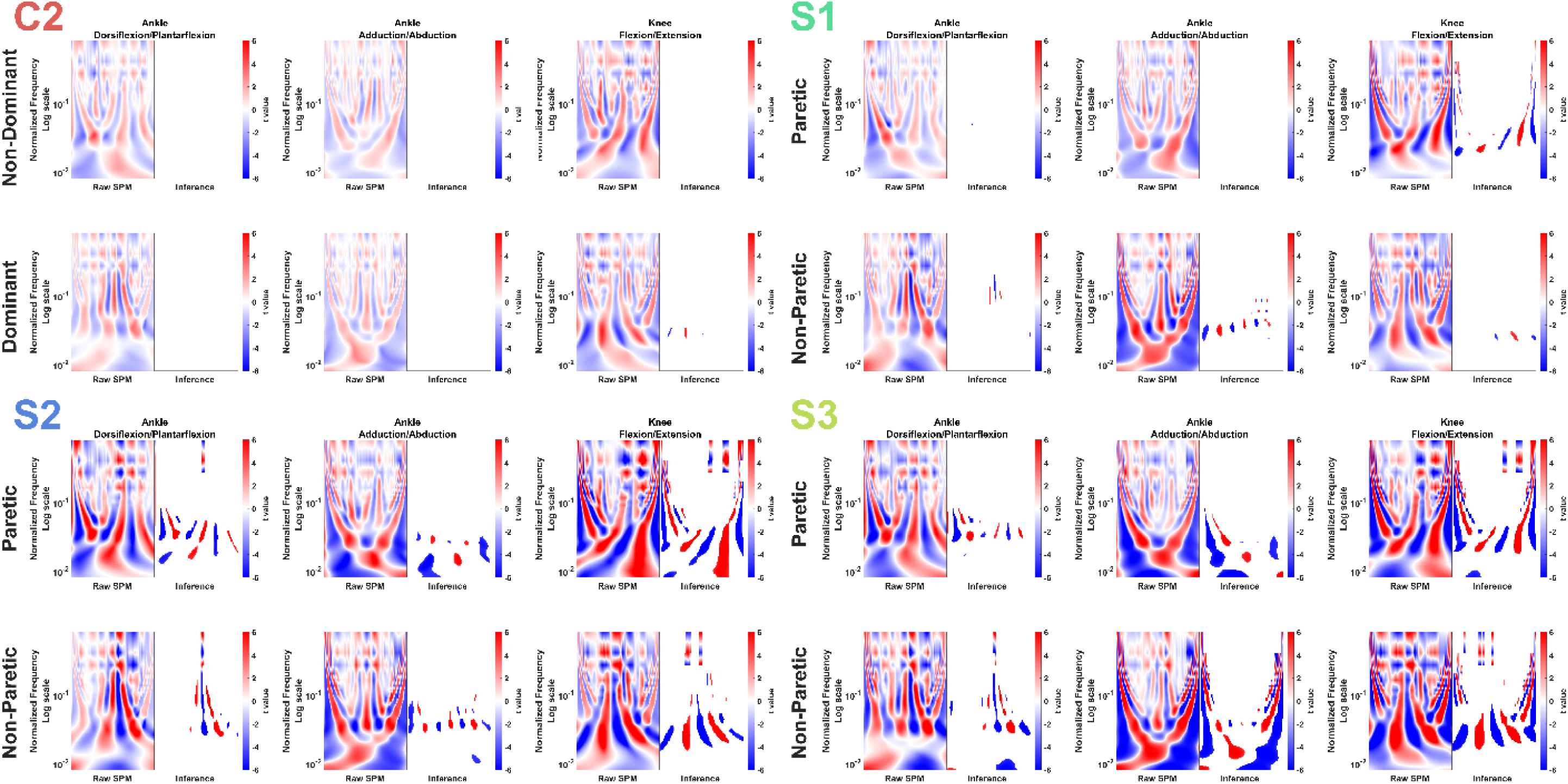
2D-Statistical Parametric Mapping t-test of the continuous wavelet transform coefficient matrix between the reference control cluster (C1) and the other clusters (Controls: C2, Stroke: S1, S2, and S3). The non-dominant side for control individuals is compared to the paretic side of post-stroke individuals, and dominant to paretic. The zones displayed in each inference panel indicate significant differences.

